# A subject-specific reversible folding model reveals geometry-driven white-matter organization

**DOI:** 10.64898/2025.12.10.693407

**Authors:** Besm Osman, Ruben Vink, Andrei Jalba, Kurt G Schilling, Maxime Chamberland

## Abstract

Tractography currently relies almost exclusively on diffusion data and seed maps, which alone remain insufficient for anatomically accurate fiber reconstruction. During brain development, white-matter fibers elongate alongside cortical folding, suggesting a close mechanistic link between cortical geometry and fiber organization. To investigate this link, we introduce a subject-specific cortical folding simulation framework that reconstructs an individual’s folding trajectory using only a structural T1-weighted image. The quasi-static, constraint-based model reverses folding to generate an unfolded, fetal-like configuration and then refolds to map the resulting volumetric deformation onto fiber organization. Validation with longitudinal fetal MRI shows that simulated folds follow biologically plausible developmental paths. Applied to adult data, deformation of simple radial fibers reproduces diffusion-derived orientation patterns across the white matter and achieves high regional correspondence. The deformation model also generates characteristic short-range U-fibers as well as long-range association and commissural pathways, all without any diffusion input or machine learning. This approach provides a new, anatomically grounded source of subject-specific fiber orientation derived solely from cortical geometry, opening avenues for geometry-informed tractography.

## 1 Introduction

Tracking fibers using diffusion MRI is currently the only viable in vivo method to estimate white matter pathways. Despite substantial methodological advances, state-of-the-art tractography still produces many anatomically implausible streamlines, overestimates the number of bundles, and explains only a fraction of known anatomy. Invalid bundles can outnumber valid ones, and current methods capture only about one-third of the true bundle extent when compared to ground truth data. (Maier-Hein et al., 2017; Rheault et al., 2019; Schilling et al., 2019). A core challenge is tracking fibers within voxels containing multiple orientations, a problem that cannot be solved by improved spatial resolution of diffusion MRI alone (Schilling et al., 2017a). Studies show that reconstructions based solely on local orientation and seeding are fundamentally limited (Schilling et al., 2019), motivating the integration of additional anatomical information.

Several strategies have demonstrated that incorporating prior anatomical knowledge improves tracking. Bundle-specific shape and curvature constraints can regularize streamline trajectories (Chamberland et al., 2017; Rheault et al., 2019), and cortical surface information has been used to mitigate gyral bias and refine streamline terminations near the gray–white matter boundary (Schilling et al., 2017b; Shastin et al., 2022; St-Onge et al., 2018; Wu et al., 2020). However, few anatomical features are both predictive of underlying fiber organization and directly obtainable from standard MRI. As a result, diffusion data remains the primary, yet incomplete, driver of tractography.

Cortical folding, or gyrification, describes the emergence of the wrinkled geometry of the cortex during fetal development, a process closely linked to both brain function and pathology (Mangin et al., 2019; Van Essen, 2020). Abnormal folding has been associated with epilepsy, autism, and schizophrenia (Garcia et al., 2018; Nordahl et al., 2007; Voets et al., 2011). Although extensively studied, the mechanisms driving cortical folding remain debated (Lewitus et al., 2013). Early theories posited that skull constraints induce cortical folding; however, experiments showed folding occurs even without skull interactions (Barron, 1950). Mechanical tension along axons may contribute to folding while maintaining compact neural circuitry (Essen, 1997; Van Essen, 2020), but since tension generally follows axonal directions rather than acting across gyri, axonal tension alone does not appear sufficient (Garcia et al., 2018). Stress-induced fiber growth may play a key role in shaping cortical morphology (Balouchzadeh et al., 2023). Recent literature focuses on differential growth between cortical layers: faster expansion of superficial tissue relative to subcortical tissue induces mechanical instability, producing folds. This phenomenon can be modeled using a thin, rapidly expanding elastic layer attached to a slower-expanding foundation, generating folding via mechanical instability and buckling.

There is a clear link between cortical geometry and axonal organization (Figure 1). Animal studies demonstrate that axons elongate in response to mechanical tension (Smith, 2009), and human imaging shows strong coupling between cortical surface phenotype and underlying white-matter architecture (Melbourne et al., 2014). Computational folding models in simplified two-dimensional settings can even predict white-matter organization, including local fiber directionality (Garcia et al., 2021). However, such models have not been exploited in tractography. While cortical surface information is used to improve seeding and orientation near the surface (Shastin et al., 2022; St-Onge et al., 2018), the *volumetric* deformations underlying fold development, potentially informative of deeper white matter fibers from structural data alone, remain unutilized.

**Figure 1:**
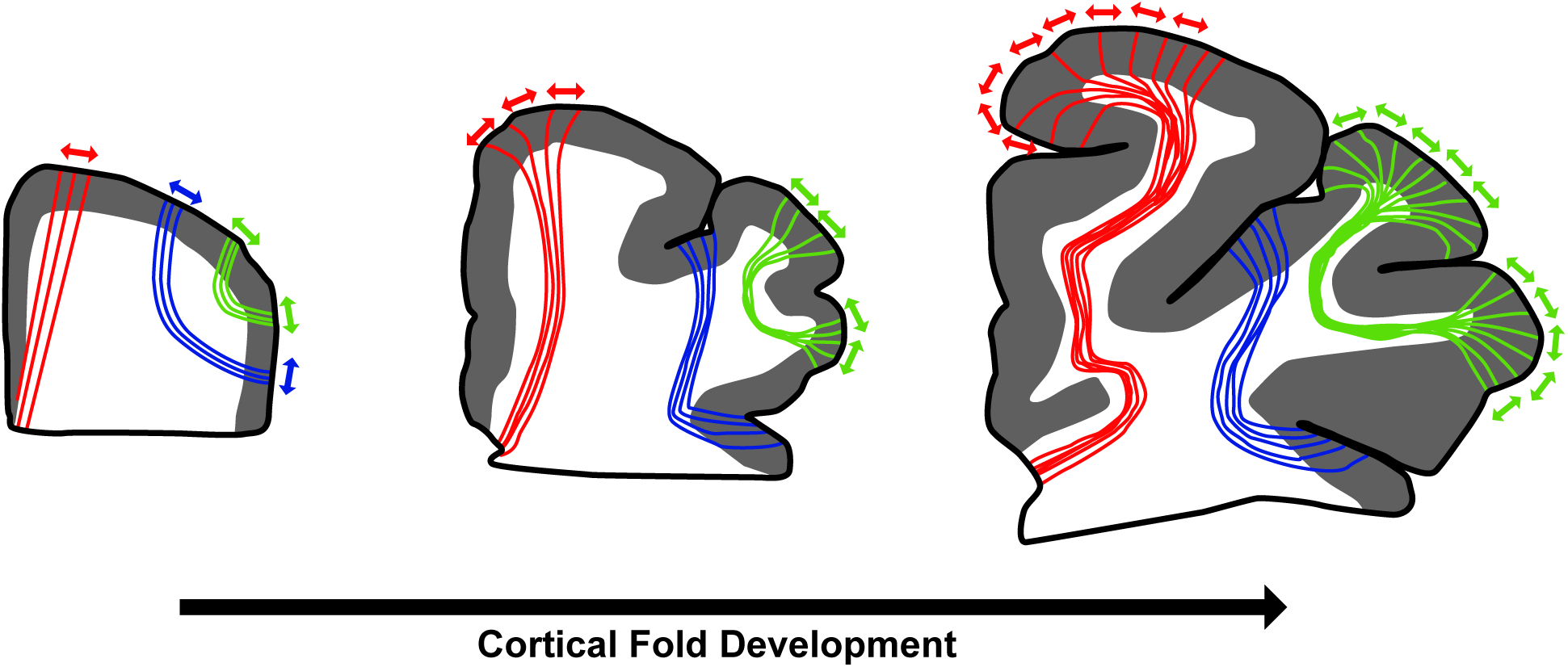
Illustration of the brain across developmental stages, showing how cortical gyrification transforms initially superficial fibers (green, blue) and deeper long-range fibers (red) into the mature folded configuration. As the cortex folds, superficial white-matter trajectories follow gyral and sulcal geometry, while deep pathways remain relatively stable, highlighting the potential mechanistic link between cortical folding and fiber organization

Existing folding simulations are limited by their inability to reproduce subject-specific patterns. Current volumetric methods use simplified two-dimensional (2D) or three-dimensional (3D) geometries, producing generic rather than subject-specific configurations (Garcia et al., 2021; Holland et al., 2015; Zhang et al., 2016). Even state-of-the-art physical models Tallinen et al., 2016 simulates growth from a 3D fetal shape to generate a cortical folding pattern resembling a human brain. Yet, when evaluated on fetal datasets, simulated folds differ from ground truth MRI data (Alenyà et al., 2022). Moreover, models linking cortical folding to white matter organization often rely on 2D simulations (Garcia et al., 2021; Solhtalab et al., 2025), making them impractical for subject-specific tractography. Interestingly, Tallinen et al., 2016 showed that repeated folding and unfolding of physical gels yields nearly identical deformation patterns, suggesting that cortical folding behaves quasi-reversibly under slow deformation.

A key missing piece is a framework that (i) captures subject-specific cortical folding patterns volumetrically, (ii) approximates their developmental trajectories, and (iii) links these deformations to white-matter orientation in a way that is practical for tractography.

In this work, we introduce a subject-specific cortical folding simulation that treats cortical development as a quasi-static, approximately reversible process. Starting from an individual’s folded brain, we “unfold” the cortex to a smooth, fetal-like configuration using a constraint-based volumetric model driven only by structural MRI, and then use the resulting deformation map to predict how simple fetal-like fiber configurations would be refolded into the adult anatomy. We first validate the unfolding–refolding behavior using longitudinal fetal MRI, and then apply the method to adult data to test whether deformation of radial fibers recovers diffusion-derived orientation patterns. This establishes cortical geometry as a direct, subject-specific anatomical prior on white-matter organization, independent of diffusion measurements. Similar approaches have been applied in the past, for example in the hippocampus, where researchers unfold the curled structure into sheet-like coordinates to facilitate tractography (Hussain et al., 2020; Karat et al., 2023) and subfield mapping (DeKraker et al., 2018). This motivates the general idea that geometry and unfolding may simplify complex folding so as to recover hidden structural pathways. Our framework requires only a single structural T1-weighted image, providing an independent anatomical source for estimating fiber orientation and opening new avenues for anatomically informed tractography.

To capture subject-specific folding dynamics, we draw on the concept of quasi-static reversibility from thermodynamics: if a system deforms slowly enough, its deformation path can be approximately retraced in reverse. An intuitive analogy is a flexible rope that bends when pushed from both ends; while predicting its forward deformation is complex, slowly pulling the ends apart allows the rope to return along nearly the same path (Figure 2).

**Figure 2:**
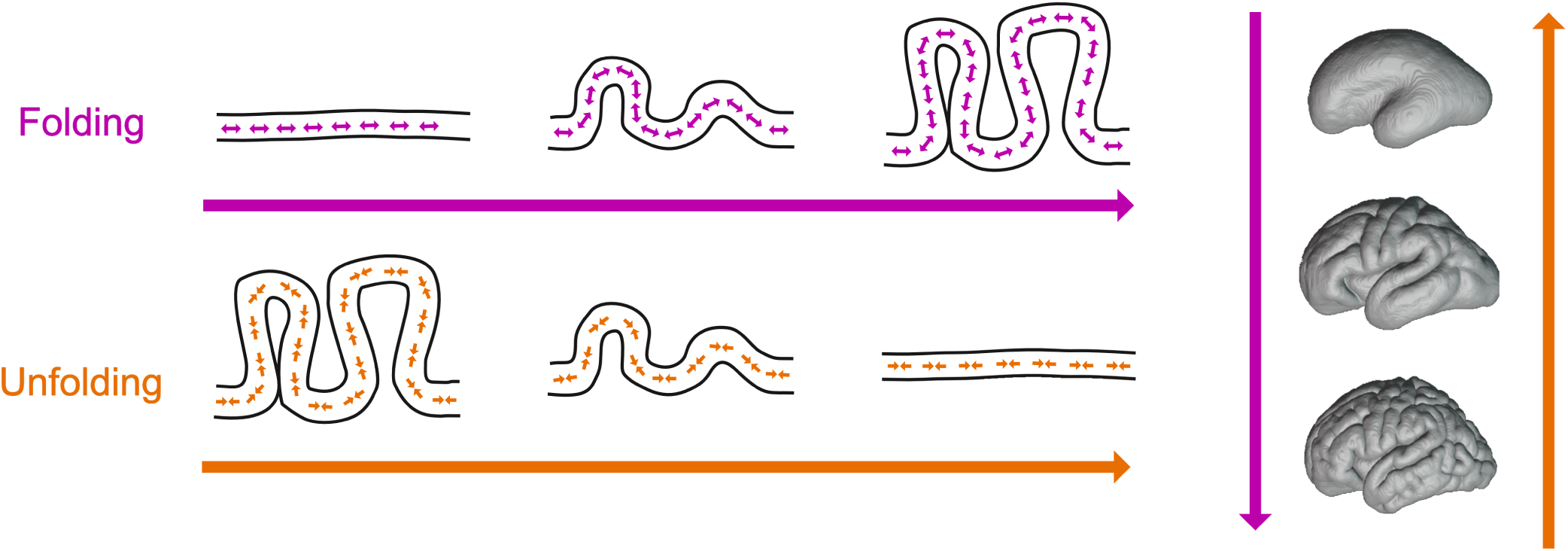
Illustration of the proposed unfolding technique. While typical cortical folding models simulate tangential expansion near the surface of a fetal-like model, our approach starts with a model derived from a postnatal scan and applies tangential contraction near the surface to approximate the cortical folding deformation in reverse.

Applying this principle to cortical development, we model folding as a quasi-static process governed by differential tangential expansion (Bayly et al., 2014). Rather than simulating growth forward from a fetal template, we begin with a subject’s folded adult (or neonatal) brain and reverse the surface constraints to approximate the deformation trajectory back to a smooth, fetal-like configuration. Each intermediate state represents a mechanical equilibrium between growth-induced stresses and tissue elasticity, allowing us to infer the deformation mapping that links unfolded and folded geometries. Here, we implement this quasi-static, subject-specific folding model and describe its components in Section 2. We then validate the framework using longitudinal fetal MRI and demonstrate its ability to predict adult white matter orientation and tractography from cortical geometry in Section 3. By establishing a direct link between cortical and white matter morphology, we aim to provide a new anatomically grounded approach for inferring white matter organization from structural imaging alone.

## 2 Methods

We developed a volumetric, constraint-based simulation pipeline (Figure 3) that (i) generates a subject-specific mesh from structural MRI, (ii) performs quasi-static unfolding that models the developmental trajectory of cortical folds, and (iii) uses the resulting deformation model to propagate idealized fibers from unfolded to folded space. Below, we describe each component and how it connects to the overall goal of linking cortical folding to fiber organization.

**Figure 3:**
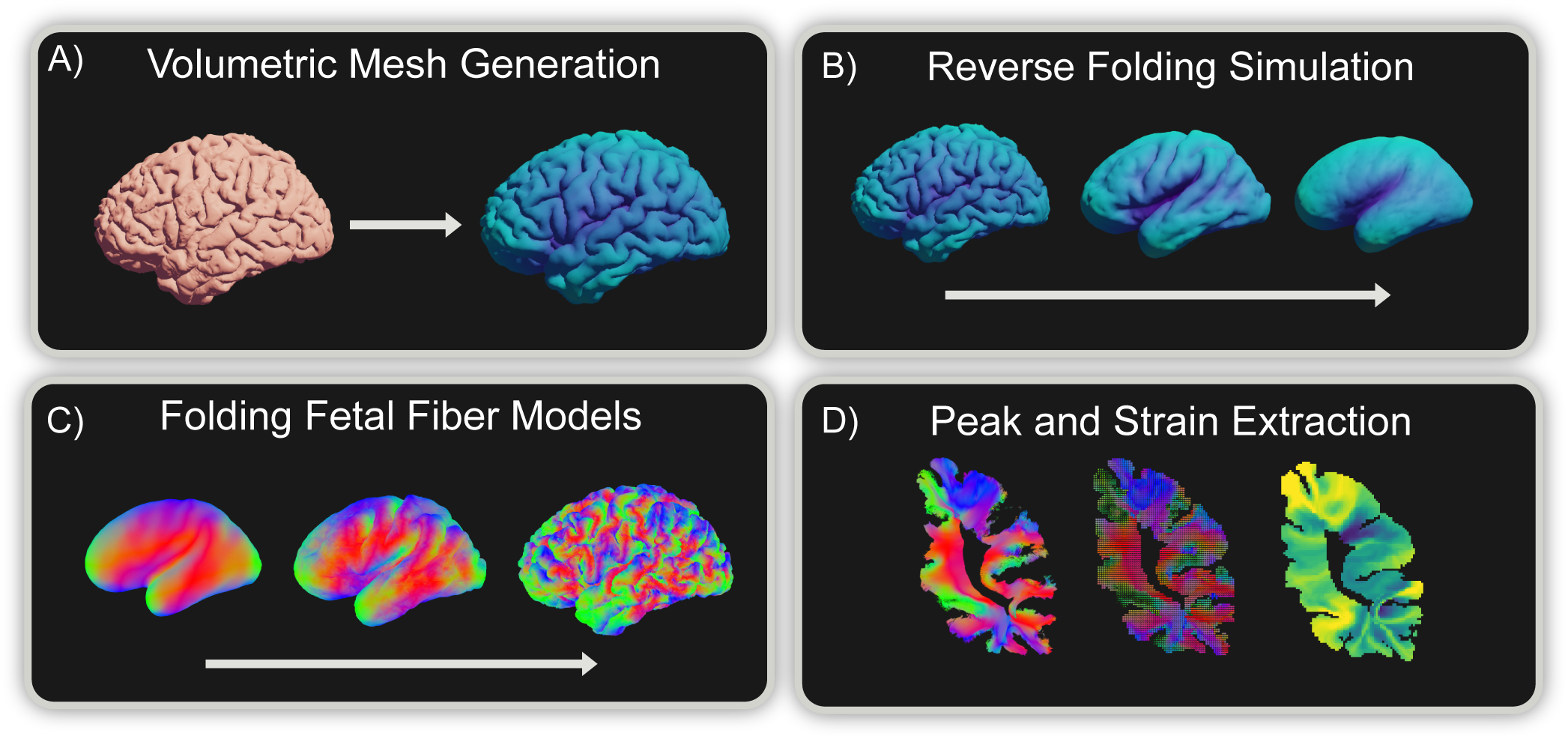
Illustration of the computational pipeline. (A) First, a volumetric mesh model is generated from the subject’s surface file extracted from either T1w or T2w MRI. (B) Next, we simulate cortical folding deformation by reversing constraints in a quasi-static simulation. From the resulting deformation mapping, we deform (C) generated fetal fiber models from the unfolded model to the folded space. The resulting fiber peaks are then extracted (D) with a map of strain and compared with the subject’s diffusion data.

### 2.1 Volumetric Meshing

Our simulation method requires a volumetric mesh that accurately represents the cortical surface from structural MRI data. It is critical that the volumetric mesh preserves the correct connectivity between distinct cortical folds to enable realistic deformation modeling. While various established tools can extract cortical surface meshes from structural MRI scans (Fischl, 2012; Shattuck & Leahy, 2002), subsequent volumetric meshing presents significant challenges.

Unconstrained meshing software (Si, 2015) connects nearby but anatomically separate folds, rendering deformation simulations highly inaccurate. Constrained meshing software typically fails when applied to surfaces extracted from automated cortical processing pipelines due to geometric irregularities such as holes and self-intersections requiring manual cleanup.

To address these limitations, we leverage a custom tetrahedralization algorithm specifically designed for cortical surfaces. The approach is robust against self-intersections while maintaining proper fold separation. The method accommodates both per-hemisphere surfaces (as generated by FreeSurfer) and complete cortical surfaces (as produced by BrainSuite). Detailed specifications of our meshing algorithm are provided in Osman et al., 2025. The subsequent pipeline components are compatible with any tetrahedral meshing technique that preserves correct connectivity and maintains anatomical separation between cortical folds.

During mesh generation, each node receives either white or gray matter designation, creating a two-tissue volumetric model. Anatomical labeling utilizes a generated Desikan atlas (Desikan et al., 2006) to assign parcellation labels to mesh nodes. Because the meshing procedure does not ensure a one-to-one correspondence between mesh surface nodes and the original cortical surface, surface node labels are assigned according to geometric proximity. Nodes positioned near opposing cortical folds may receive incorrect labels, which are automatically relabeled by an iterative neighbor-based majority voting step across the surface mesh, smoothing local inconsistencies and enforcing contiguous regions with coherent labels.

### 2.2 Mechanical Model

Cortical folding deformation is simulated with a constraint-based formulation implemented in extended position-based dynamics (Macklin et al., 2016). The volumetric mesh is modeled using a neo-Hookean material energy model

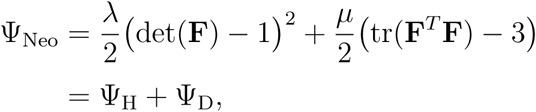

where **F** is the deformation gradient, and *λ* and *µ* are the Lamé parameters. The energy is maintained through stable tetrahedral constraints (Macklin & Muller, 2021).

Brain tissue exhibits viscoelastic behavior, demonstrating both viscous and elastic properties. To incorporate viscous behavior within our model, we adjust the rest pose matrix used to determine the deformation gradient **F** toward the current tetrahedral pose, allowing deviatoric energy to incorporate pose changes and thereby modeling viscoelastic deformation. Since our primary concern is mapping the deformation caused by folding, we ignore the isometric growth of the brain during development, as it does not contribute to the deformation mapping.

To model tangential expansion forces, we introduce surface area constraints per face on the surface of the volumetric mesh:

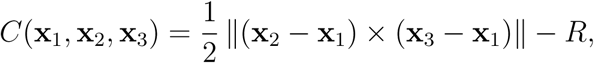

where **x**_1_, **x**_2_, **x**_3_ are the face vertices and *R* is the rest area that starts as the face area and decreases over time. The locally decreasing rest area pushes nodes toward the direction of minimizing surface area, while the neo-Hookean hydrostatic energy counteracts volumetric change, and the relaxed deviatoric energy reduces shape deformation.

XPBD minimizes local constraint violations, thus may not find the globally optimal configuration. Surface area constraints in highly curved regions can be problematic, as a smooth surface may require an increase in hydrostatic constraint violation near sulci to reduce overall area constraint violation. Since global constraint solving for high-quality models is impractically slow, we instead bias our constraint solving toward smooth surfaces by introducing a smoothing constraint per surface area neighbor pair to converge toward global area reduction, i.e.,

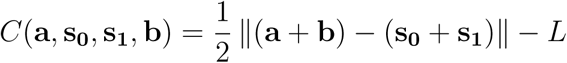

where **s_0_**, **s_1_** are the shared edge nodes of two adjacent triangles, **a**, **b** are the remaining nodes, and *L* is the rest distance. This constraint increases smoothing by moving the shared edge center toward the center of **a**, **b** by decreasing *L* over time. Since, under fixed volume, a sphere minimizes surface area (isoperimetric inequality), this formulation provides an efficient local approximation to global surface-area reduction.

### 2.3 Simulation Parameters

Material properties were chosen within experimentally reported ranges for brain tissue: a Poisson ratio of *ν* = 0.48 (Morin et al., 2017) and tissue stiffness values of 1.895 kPa for white matter and 1.389 kPa for gray matter (Budday et al., 2015). For mass computation, a tissue density of 1040 kg/m^3^ is distributed across nodes based on tetrahedral volume, since both white and gray matter exhibit approximately equal density (Abdi et al., 2024; Yu et al., 2018). The viscoelastic factor is determined through experiments with a simulated 1 mm^3^ white matter cube model to achieve behavior consistent with tissue viscoelasticity described in the literature (Budday et al., 2020), scaled to the chosen timescale.

To maintain quasi-static simulation conditions within a reduced timeframe, we set viscous damping to ∞ such that acceleration remains zero regardless of timescale. Consequently, the resulting deformation is purely constraint-driven, resolving to equilibrium at each timestep given sufficient XPBD substeps.

Remaining XPBD parameters were tuned empirically for smooth convergence and biological plausibility by comparison with a fetal developmental atlas (Xia et al., 2019). These parameters do not represent precise biophysical constants but control numerical stability and relative stiffness between tissue layers.

### 2.4 Deformation Map

The resulting simulation provides a time-evolving mesh with changing node positions and constant connectivity structure. For subsequent mapping of fibers between folded and unfolded states, we define a deformation function Φ*_t_* : ℝ^3^ → ℝ^3^ that maps any point **p** contained in the volumetric model between unfolded and folded configurations,

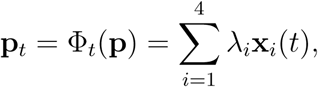

where **x***_i_*(*t*) are the tetrahedral vertex positions at time *t* ∈ [0, 1], where *t* = 0 is the unfolded configuration and *t* = 1 is the folded configuration. *λ_i_* are the barycentric coordinates of the point in the reference configuration *t* = 0.

Time-varying constraints can produce deformations of differing magnitudes across simulation timesteps. To ensure uniform temporal sampling of *t*, we normalize the deformation sequence based on node movement. We measure the Euclidean displacement of nodes between each pair of consecutive simulation states, then adjust the temporal spacing so that periods of greater physical change are increased in duration. This creates a normalized parameter *t* where equal time steps correspond to roughly equal amounts of physical deformation.

### 2.5 Fiber Modeling

Using the deformation function Φ*_t_*(**p**), various fetal-like fiber geometries can be deformed from the unfolded state to any time *t* ∈ [0, 1]. A deformation bundle is defined as a set of initial fibers in the unfolded space with the deformation function from the corresponding simulation run. Thus, bundles, unlike diffusion-based bundles, can be computed and visualized at intermediate *t*, where *t* = 1 represents the fibers deformed to the folded configuration.

For validation, we employ a simple radial fiber model, in which straight fibers originate at the cortical surface and extend inward along local surface normals. A volumetric field of inward-facing normals is computed by voxelizing the unfolded surface and averaging neighboring vectors, followed by smoothing and normalization to reduce discretization artifacts. As folding progresses, the fibers are advected by the deformation field and bend, twist, or stretch according to local strain. This produces a predicted fiber orientation field derived purely from geometry, without any diffusion input or machine learning.

We also demonstrate that coherent fiber bundles can be synthesized using the same deformation-based framework. Bundle endpoints were obtained from TractSeg (Wasserthal et al., 2018). They were then connected in the unfolded configuration using randomized Bézier curves defined by the two atlas endpoints and a single control point specifying the curve midpoint, akin to stem-based approaches (Calixto et al., 2025; Hau et al., 2016, 2017). These synthetic fibers were then subjected to the folding deformation to generate corresponding tract bundles in the folded space.

### 2.6 Dataset description

The validation dataset for the unfolding simulation consists of longitudinal structural MRI from the Developing Human Connectome Project (dHCP) (Edwards et al., 2022), containing fetal 33GA and neonatal 42GA scans. At 33GA, primary and secondary folds are visible. By the neonatal scan, tertiary folding has progressed.

Image acquisition methods and parameters are detailed in (Edwards et al., 2022). Imaging was carried out on a 3T Philips Achieva scanner using a dedicated neonatal 32 channel phased array head coil and customized patient handling system (Rapid Biomedical GmbH, Rimpar, Germany) (Hughes et al., 2017). T1w multi-slice FSE images (TR/TI/TE = 4795/1740/8.7 ms, SENSE factor 2.27 (axial) and 2.66 (sagittal)) were acquired in sagittal and axial slice stacks with an in-plane resolution of 0.8x0.8mm^2^, and fused into a single 3D volume at 0.8mm isotropic volume (Cordero-Grande et al., 2016). Diffusion data were acquired with 4 shells (b = 0 s/mm2: 20, b = 400 s/mm2: 64, b = 1000 s/mm2: 88, b = 2600 s/mm2: 128) (Hutter et al., 2018) (MB factor 4, SENSE factor 1.2, partial Fourier factor 0.86, in-plane resolution 1.5 × 1.5 mm, 3 mm slices with 1.5 mm overlap, TE = 90 ms, TR = 3800 ms) and preprocessed to correct for motion, eddy currents, and susceptibility distortion (see (Hughes et al., 2017) for details). Cortical surfaces were genereated from the T1-weighted image using infant FreeSurfer (Zöllei et al., 2020).

For validating deformation fibers, adult structural and diffusion data were acquired from the Human Connectome Project Young Adult dataset (Van Essen et al., 2013). Cortical surfaces were generated from the T1-weighted image using BrainSuite cortical surface extraction (Shattuck & Leahy, 2002) resulting in a surface with 585k vertices.

### 2.7 Evaluation

The framework assumes cortical folding is approximately reversible, such that reverse simulation reproduces trajectories consistent with forward developmental folding. We therefore validated the model at two levels: 1) developmental plausibility, and 2) fiber-orientation prediction. For 1), longitudinal structural MRI from the Developing Human Connectome Project (dHCP) was used, containing a fetal scan at 33 gestational weeks (GA) and a neonatal scan at 42 GA for the same subject. Reverse simulations were run from the 42 GA surface until the gyrification index matched that of the 33 GA scan, and the resulting surfaces were compared to evaluate whether simulated folds followed the correct developmental elongation and direction. For 2), we used an adult subject from the Human Connectome Project (HCP) (Van Essen et al., 2013). The pipeline was applied to the adult structural image to produce an unfolded, fetal-like surface. The generated volumetric mesh contains approximately 160,000 nodes with approximately 850,000 tetrahedra. Within the unfolded model, radial fibers were generated and subsequently deformed into the folded state using the simulation mapping. Primary peaks for validation were extracted from orientation distribution functions generated using MRtrix’s multi-shell multi-tissue constrained spherical deconvolution (Jeurissen et al., 2014; Tournier et al., 2019). Cosine similarity between simulated and diffusion-derived primary peaks was calculated voxel-wise in white matter (excluding interhemispheric regions) as *C*(**v**) =| **a**(**v**) · **b**(**v**) |, where **v** is a voxel coordinate, **a** is the normalized primary diffusion peak, and **b** is the normalized simulated primary peak.

## 3 Results

We evaluated whether the proposed quasi-static reverse-folding framework (i) reproduces realistic cortical developmental trajectories and (ii) predicts white-matter orientations consistent with diffusion MRI, despite relying solely on structural data. Results are organized as follows: Section 3.1 validates developmental plausibility, Section 3.2 characterizes the resulting strain patterns, Sections 3.3, 3.4 examine how radial fibers deform and align with diffusion peaks, and Section 3.5 illustrates tractography-relevant fiber geometries.

### 3.1 Structural Alignment

To test whether reverse folding produces biologically plausible trajectories, we compared simulated cortical surfaces to longitudinal fetal MRI from a single subject scanned at 33 and 42 gestational weeks (GA) from the Developing Human Connectome Project (dHCP). The simulation was initialized from the neonatal (42 GA) surface and evolved backward until its gyrification index matched that of the 33 GA scan.

Figure 4 compares the simulated fetal-like surface to ground-truth data. The reverse simulation closely reproduces the direction and extent of fold elongation: secondary and tertiary folds visible at 42 GA flatten in the simulated 33 GA surface, while primary sulci are preserved. Spatial correspondence between simulated and observed folding patterns is particularly strong in temporal and parietal regions, where primary folds straighten and secondary folds shorten. The left temporal pole exhibits an additional fold matching the ground truth, whereas the right hemisphere shows a slightly smoother pattern, consistent with reported inter-hemispheric asymmetries in folding timing. Overall, the model captures subject-specific trajectories of fold emergence and elongation, demonstrating that quasi-static reverse folding provides a realistic approximation of developmental cortical evolution.

**Figure 4:**
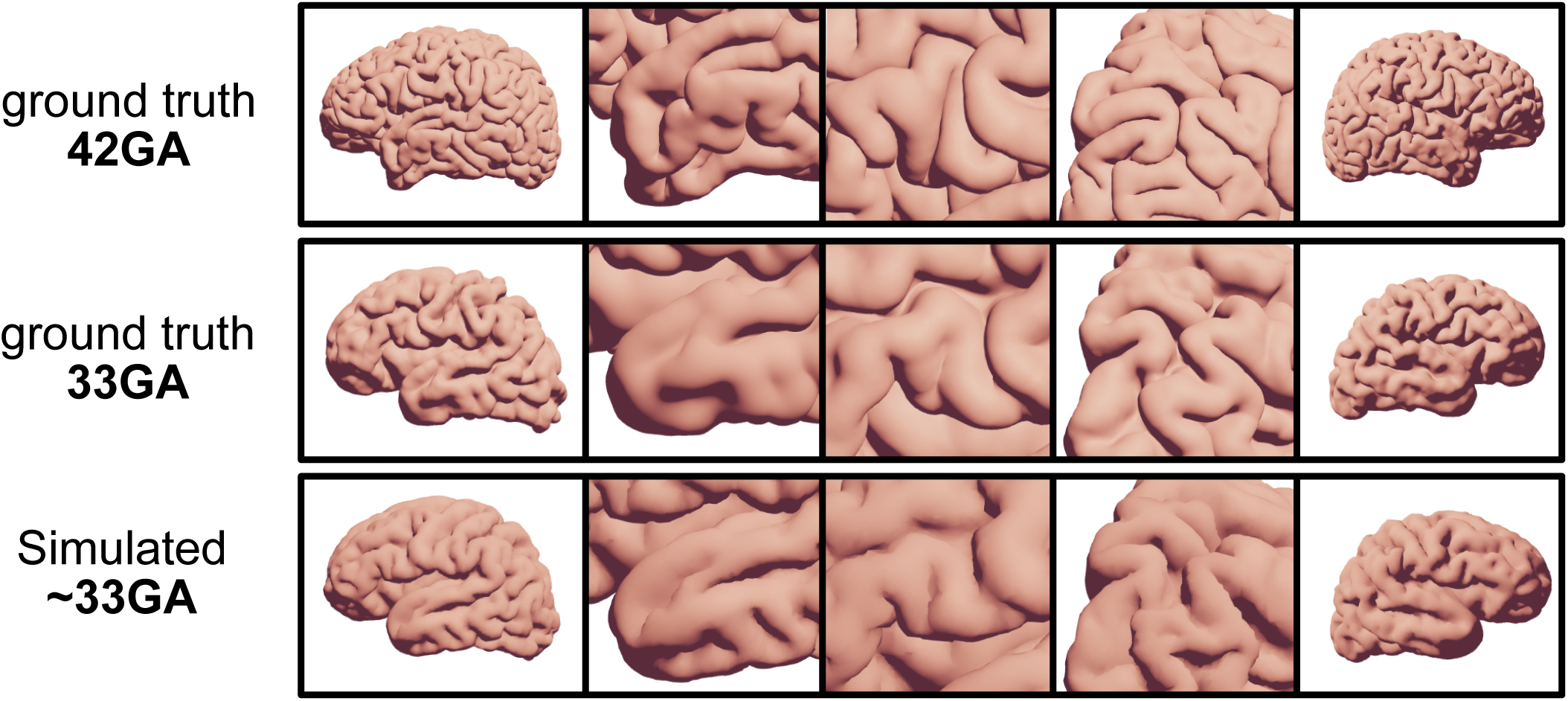
Comparison between simulated folds and fetal MRI ground truth. Top row: neonatal scan at 42GA. Middle row: fetal scan of the same subject at 33GA. Bottom row: simulated volumetric model at 33GA (matched by gyrification index). Columns highlight different regions; the first three images show the left hemisphere simulation, the last two columns right hemisphere.

### 3.2 Strain maps

We next quantified how cortical folding deforms local white matter by computing strain maps from the simulated deformation fields (Figure 5). In this context, strain measures the elongation of radial fibers during the unfolding–refolding process. Approximately ten million synthetic radial fibers were sampled throughout the cortical volume in the adult subject. For each voxel (0.325 mm isotropic), we calculated the stretch ratio as the ratio between the average folded fiber length and its original unfolded length, then defined strain as this elongation ratio minus one. Because the unfolded fiber length is constant across all samples, the strain field reflects the local fractional change in radial distance caused by cortical deformation. Positive values indicate elongation or stretching, whereas negative values correspond to local compression.

**Figure 5:**
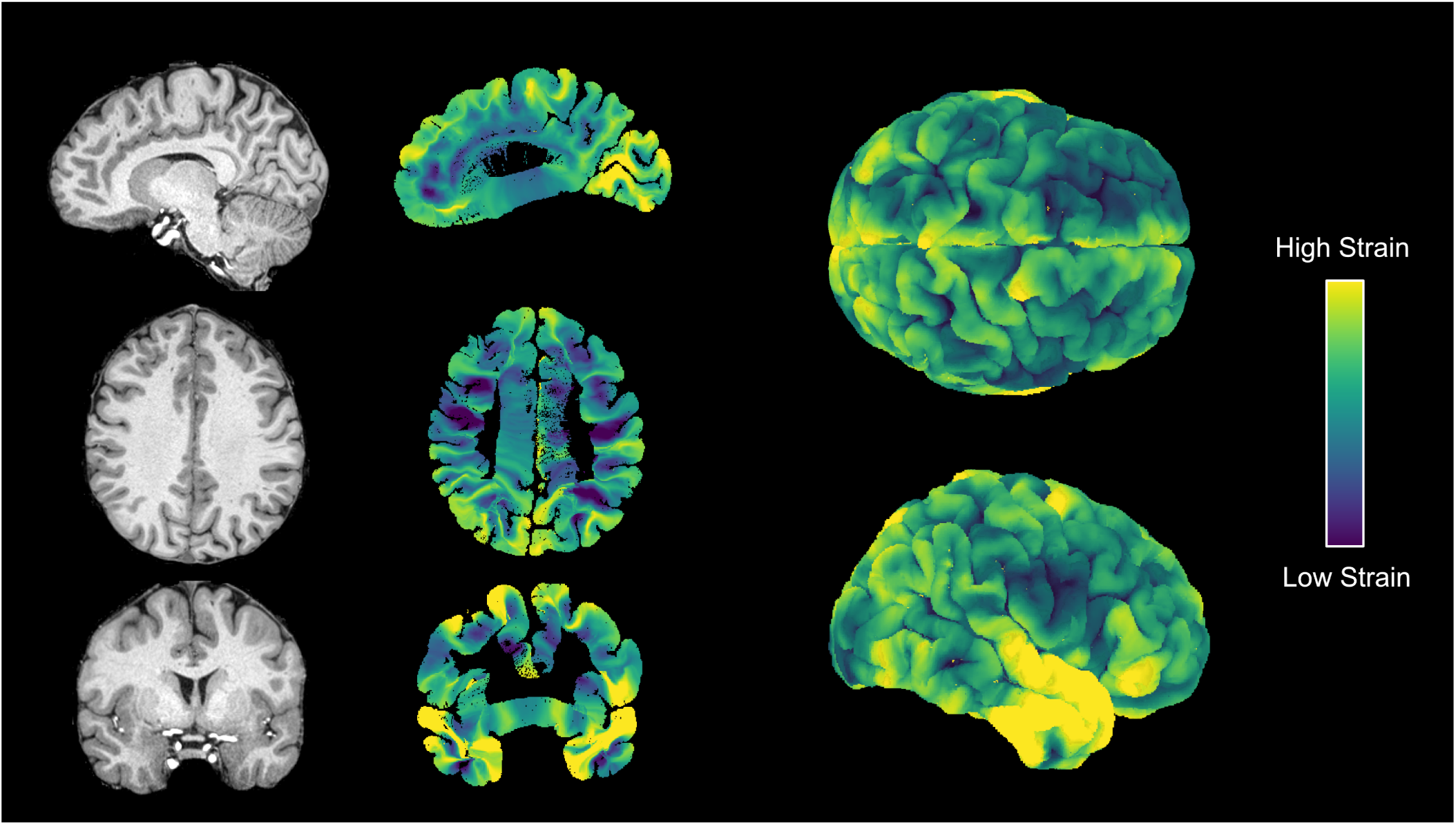
Strain maps of cortical folding. Strain represents the local elongation of white matter fibers relative to the unfolded state. Higher strain (yellow) indicates regions undergoing significant biological stretching or shearing during gyrification, whereas lower strain (blue/purple) indicates minimal deforma-tion or compression. Strain is heterogeneous across the brain, with higher strain in the temporal lobe, adjacent to the Sylvian fissure, and in U-shaped pattern immediately adjacent to sulci.

The resulting maps reveal highly heterogeneous deformation across the cortex (Llinares-Benadero & Borrell, 2019). The strongest elongation occurs around the Sylvian fissure, temporal lobe, and perisylvian regions, consistent with areas undergoing complex tertiary folding during late gestation (Im & Grant, 2019; Leroy et al., 2015; Xia et al., 2019). Strain forms distinctive U-shaped patterns along sulcal banks, capturing the mechanical stretching of tissue between adjacent gyri. These regions coincide with known concentrations of short association fibers (Van Dyken et al., 2024), suggesting that the geometry-driven deformation underlying cortical folding directly shapes the organization and potential elongation of superficial white matter. The simulation can be seen at various timepoints in Suppl. Video 1.

### 3.3 Folding Radial Fibers

To examine how cortical deformation reorganizes white-matter pathways, we generated an initial scaffold of radial fibers consisting of straight streamlines extending perpendicularly from the cortical surface into the white matter of the unfolded model and applied the simulated folding deformation.

As shown in Figure 6, originally straight radial fibers bend and twist as folding progresses, forming tangential and U-shaped trajectories near the cortical surface. In gyral crowns, fibers remain largely radial, whereas along sulcal banks they curve sharply toward opposing walls, creating arches that connect adjacent gyri. This purely geometry-driven deformation reproduces canonical superficial white-matter patterns such as gyral fanning and inter-gyral loops, demonstrating that realistic local fiber geometries can emerge from mechanical folding alone, without diffusion data (see Suppl. Videos 2, 3).

**Figure 6:**
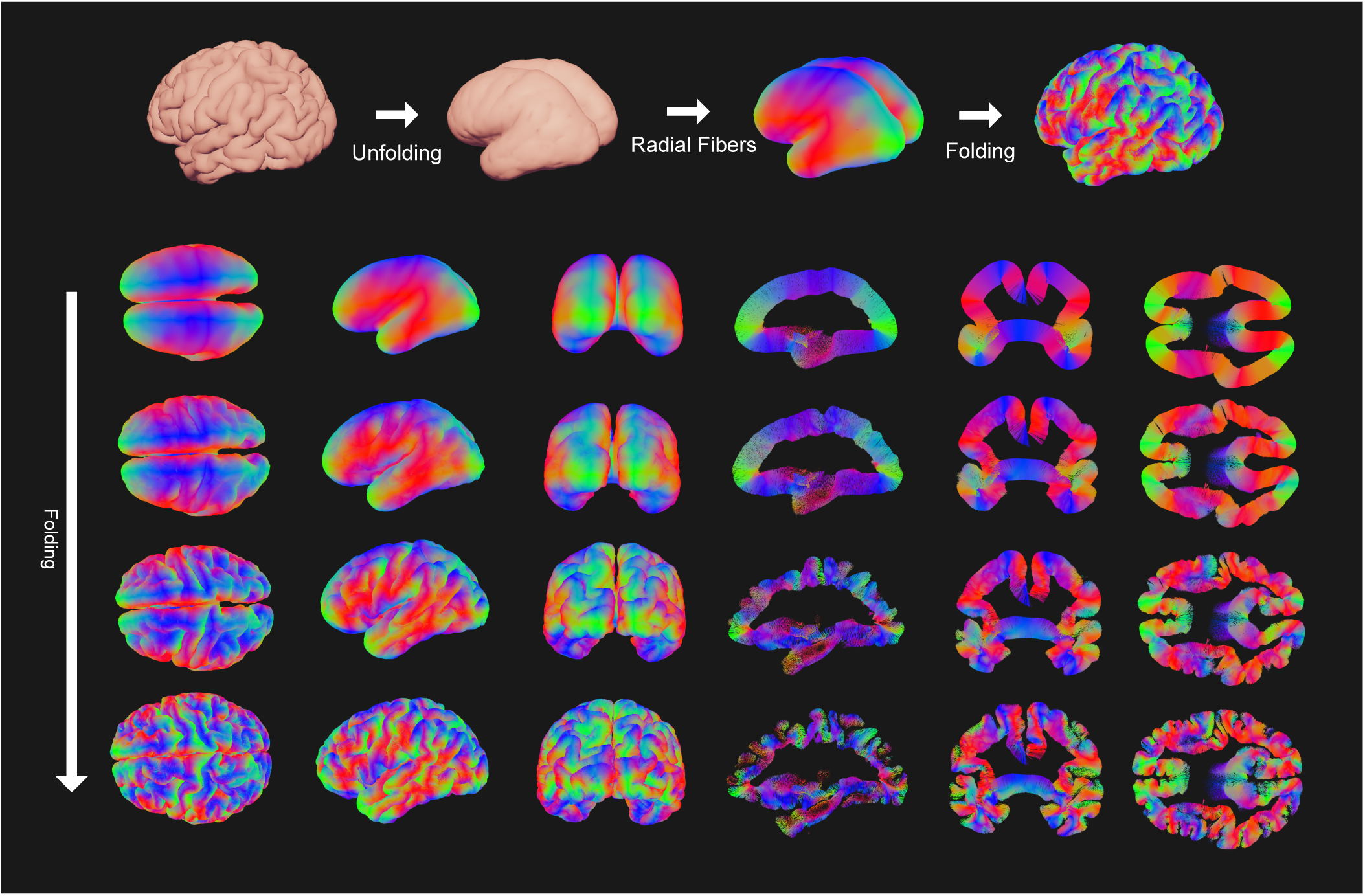
Illustration of radial fibers being folded. Top: steps starting from an adult subject, unfolding via quasi-static simulation, followed by creation of radial fibers in the unfolded model, which are then folded to the adult model. Bottom: different views and slices, each row corresponding to increasing t (top to bottom: 0, 1/3, 2/3, 1).

### 3.4 Peak Alignment

The pipeline was run (Figure 7), and cosine similarity scores were grouped by Desikan parcellation label in Figure 8. Out of 21 evaluated non-interhemispheric regions, 20 regions show a strong alignment between diffusion peaks and deformation peaks, ranging from 0.57 to 0.84, with an overall average of 0.71. The Insula was the only region with a score less than 0.5. This is consistent with this region’s flatter, less convoluted structure and divergent growth trajectory relative to other lobes (Mallela et al., 2023), making it less suitable for the folding framework. Scores by white matter depth remain relatively consistent up to 6 mm from the surface, after which alignment decreases. Peaks in gyri regions are more aligned than in sulci, and deeper white matter in gyri also matches orientation. While the model shows similar bending near gyri, radial fibers do not capture sulci bending, where the principal stress direction is orthogonal to the radial fibers. Some structures are slightly translated anteriorly relative to ground truth, producing low voxel-wise cosine similarity despite visual alignment (see Figure 1 in supplementary material).

**Figure 7:**
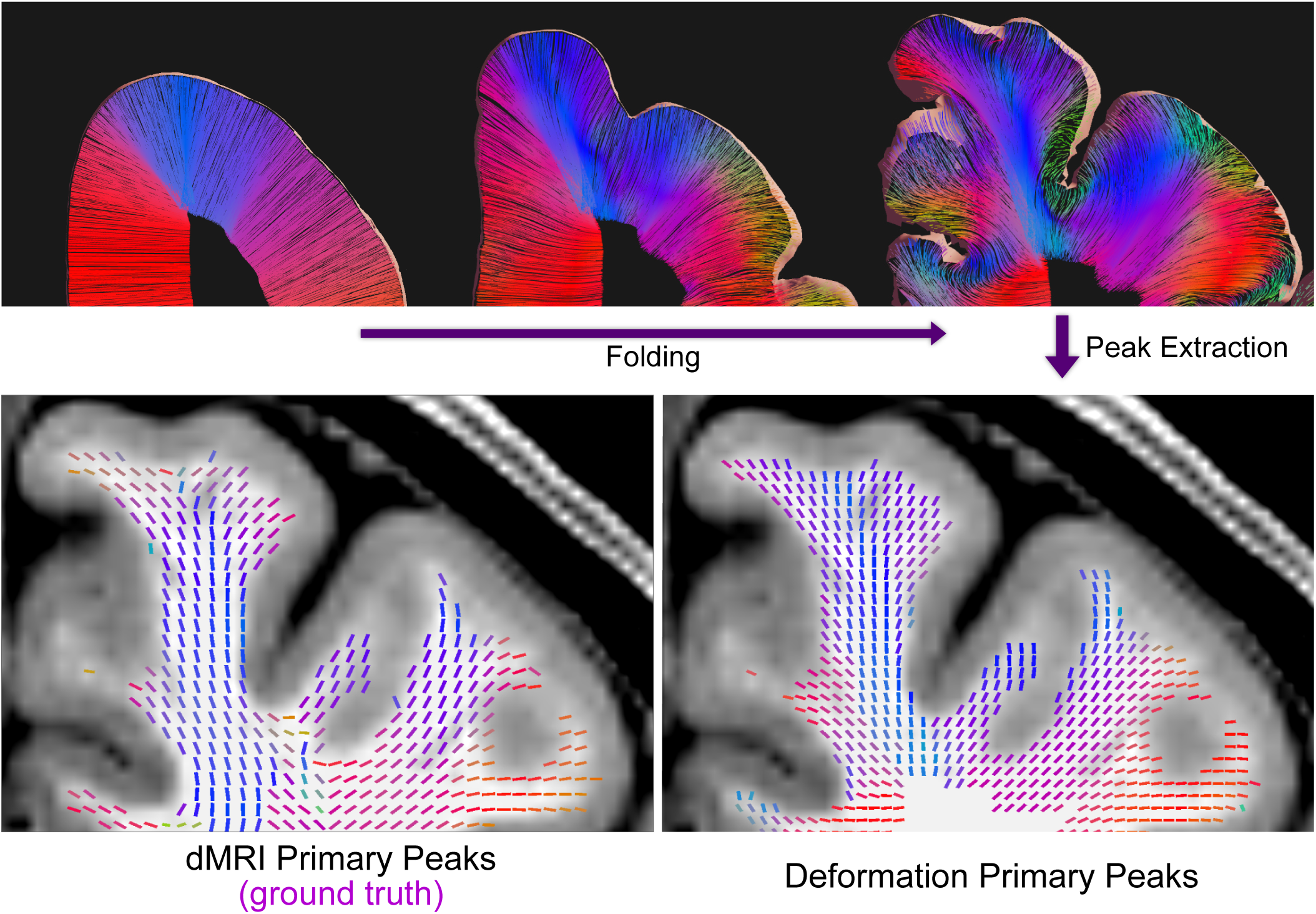
Comparison of diffusion primary peaks with peaks extracted from deformed radial fibers. Top: generated radial fibers from the unfolded to the folded model. Bottom: primary peaks of white matter voxels. Bottom left: diffusion MRI primary peak. Bottom right: fiber-derived primary peaks in folded configuration.

**Figure 8:**
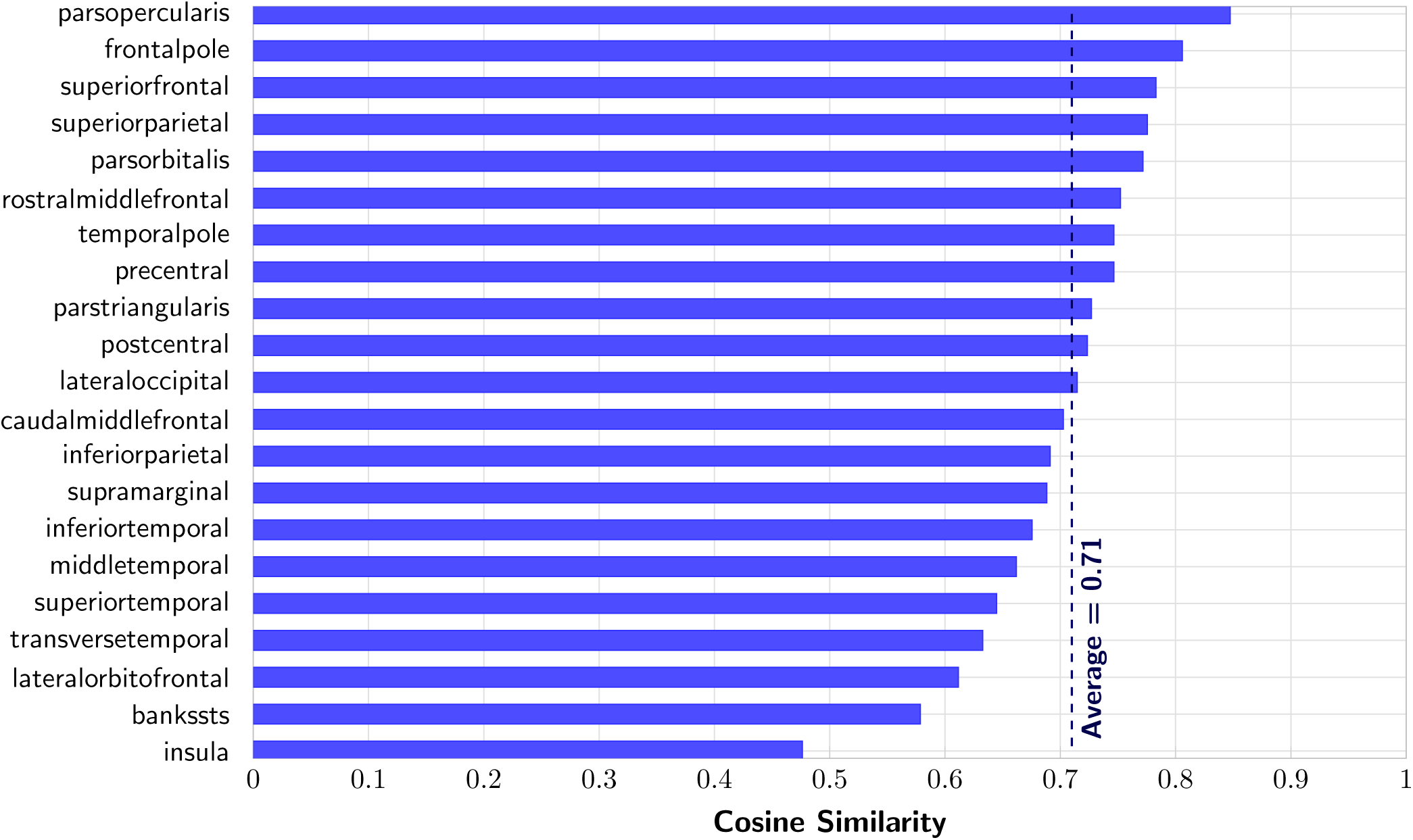
Alignment of white matter deformation peaks with diffusion primary peaks by parcellation region. The dashed line indicates the average cosine similarity of 0.71 (excluding the insula).

### 3.5 Tractography

We then evaluated whether the geometry-derived orientation fields can reproduce known short-range association pathways. Figure 9 illustrates simulated streamlines in the central sulcus, a region characterized by prominent U-shaped fibers bridging the pre- and post-central gyri (Pron et al., 2021).

**Figure 9:**
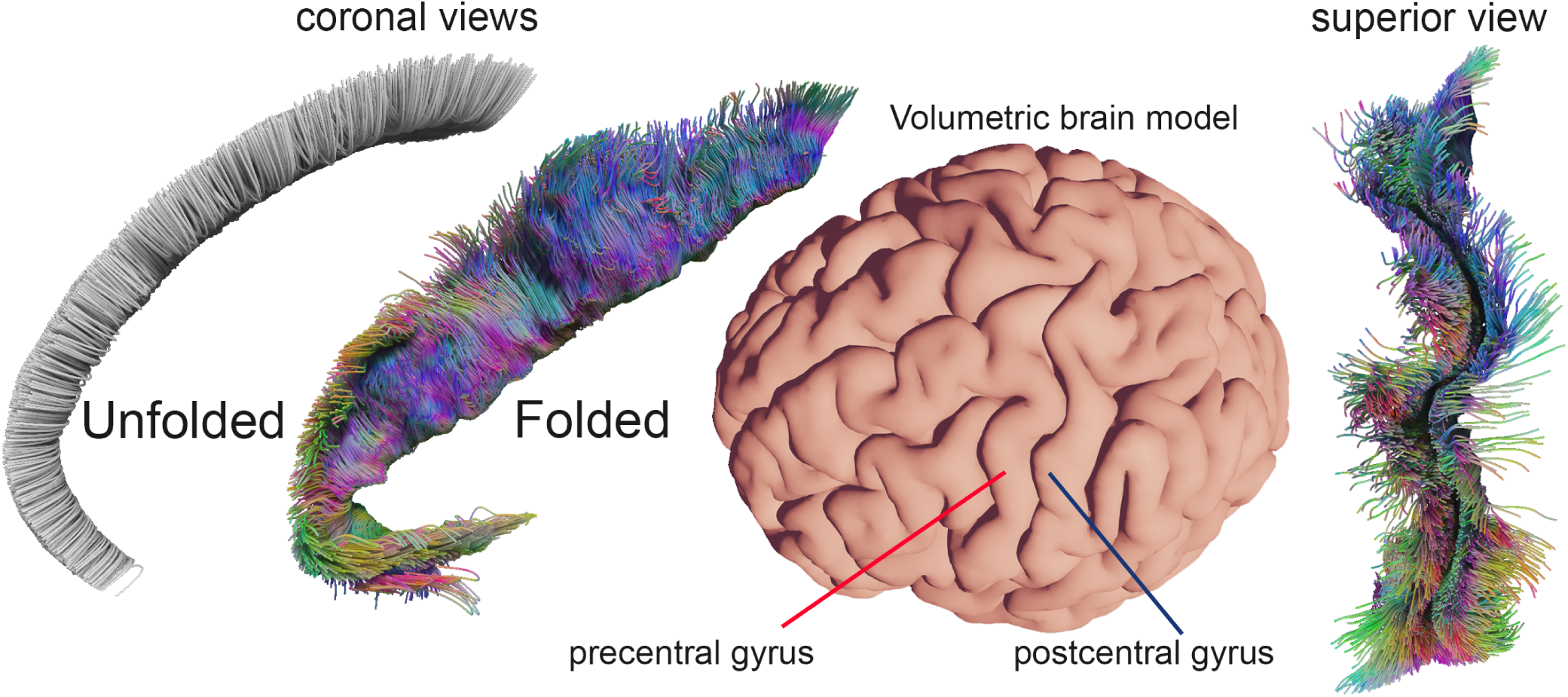
Simulated short-range U-shaped association fibers around the central sulcus, produced purely by geometry-driven deformation of fibers. Color encodes local streamline orientation, demonstrating continuous curvature along sulcal banks and smooth transitions across gyral crowns. No diffusion data were used in generating these trajectories.

Without any diffusion input, the deformation model generated a continuous sheet of short association fibers running tangentially beneath the cortical surface. Within this sheet, clusters of streamlines with subtly distinct orientations and curvatures formed along the sulcal extent, closely resembling diffusion-derived sub-bundles previously reported in this region. These trajectories emerged naturally from the deformation of initially radial fibers and remained confined to the superficial white matter, showing that subject-specific cortical geometry alone constrains the organization of local association pathways. An animated version of the result can be seen in Suppl. Video 4.

Finally, we compared the resulting deformation-derived bundles with their TractSeg (Wasserthal et al., 2018) version (Figure 10). Despite using only atlas-defined endpoints and no diffusion data or filtering, the deformation bundles reproduced the orientation and overall shape of the diffusion-derived pathways. In particular, the arcuate fasciculus, demonstrates that anatomically plausible tracts can emerge purely from geometry-driven deformation.

**Figure 10:**
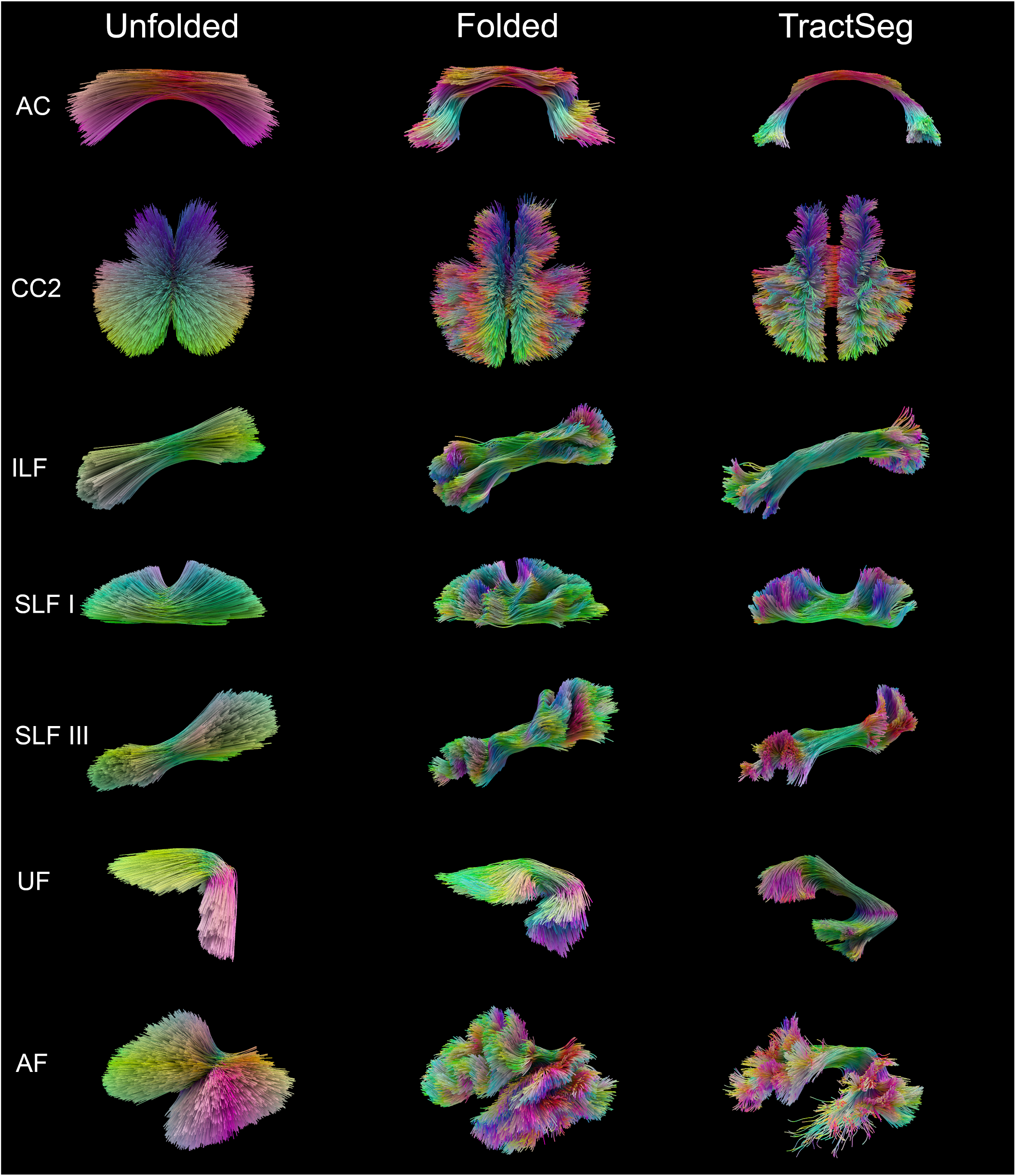
Superficial and deep white matter tractography obtained from the deformation-based model in the unfolded stage (left), folded stage (middle) and diffusion MRI tractography based on TractSeg (right) for seven representative bundles.

## 4 Discussion

This study introduces a subject-specific cortical folding simulation that reconstructs an individual’s deformation trajectory from a single structural MRI. By reversing folding constraints under quasi-static conditions, the framework produces a biologically plausible “unfolded” configuration and a volumetric deformation field that maps between folded and unfolded states. Applying this deformation to synthetic radial fibers yields orientation fields that closely match diffusion-derived white-matter patterns, even though no diffusion information is used. Together, these results demonstrate that cortical geometry encodes substantial information about white-matter organization, providing an anatomically grounded prior for tractography.

### 4.1 Linking cortical geometry and white-matter organization

Our findings support the hypothesis that cortical folding and white-matter architecture are mechanistically coupled. The close alignment between deformation-derived orientations and diffusion-based fiber peaks suggests that a large component of adult fiber organization arises from the mechanical transformations induced by folding. In the model, simple radial fibers in an unfolded, fetal-like configuration are sufficient to reproduce many of the complex curvature and fanning patterns observed in adult white matter. This result resonates with developmental evidence showing that axons elongate and reorient in response to mechanical forces generated during cortical expansion (Bayly et al., 2014; Garcia et al., 2021; Smith, 2009; Tallinen et al., 2016).

Our findings also relate to Meynert’s principle (Catani, 2025; Meynert, 1885), which states that short association fibers lie superficially beneath the cortex whereas longer pathways course progressively deeper in the white matter. In our deformation model, radial fibers undergo the strongest bending and elongation near the cortical surface—particularly along sulcal banks—naturally forming superficial U-shaped trajectories, while deeper fibers remain comparatively stable and retain long-range orientations. This depth-dependent arrangement emerges directly from geometry-driven cortical folding, suggesting that the laminar organization described by Meynert may arise, in part, from the mechanical consequences of gyrification.

While diffusion MRI has traditionally been treated as the primary source of fiber orientation, our results imply that geometry alone can constrain the feasible orientation space. The correspondence between predicted and measured fiber peaks across cortical regions (mean cosine similarity 0.7) indicates that a substantial component of the diffusion signal’s directional structure may reflect orientation driven by deformation. This provides a new interpretive link between macroscale morphology and microstructural orientation.

### 4.2 Strain and variability

The strain analysis revealed spatially heterogeneous deformation patterns, with strongest elongation along sulcal banks and perisylvian regions—areas undergoing late-developing tertiary folding. These regions are also known to exhibit higher interindividual variability and are frequently implicated in neurodevelopmental disorders such as dyslexia, autism, and schizophrenia (Im & Grant, 2019; Nordahl et al., 2007; Voets et al., 2011). The observed strain hotspots may therefore correspond to zones of mechanical stress concentration where white-matter pathways could be susceptible to variation or disruption (Zhou et al., 2025). This pattern offers a potential biomechanical perspective that may contribute to understanding developmental vulnerability. Because folding locally stretches and compresses the cortical mantle, small differences in timing or growth rates could amplify into macroscopic variability in cortical shape and connectivity. The model’s strain maps thus provide a quantitative framework for studying how differential expansion contributes to both normal variability (Croxson et al., 2018) and atypical neurodevelopment.

The deformation-based strain maps generated by our model may also be relevant for biomechanical research in traumatic brain injury (TBI). Because regions experiencing high developmental strain correspond to areas of complex folding, structural variability, and dense short-range connectivity, these locations may represent zones of increased mechanical sensitivity during head impact. Incorporating such subject-specific strain distributions into finite-element TBI simulations (Zhou et al., 2025) could improve predictions (He et al., 2024) of axonal damage risk by linking macroscopic loading patterns to locally stretched or shear-prone white-matter geometry. More broadly, geometry-derived strain fields may serve as an anatomical prior for identifying individualized vulnerability profiles, offering a potential bridge between developmental biomechanics and injury biomechanics (Zhou et al., 2021).

Moreover, the finding that superficial white-matter bundles align with high-strain regions suggests a mechanical basis for their spatial distribution. The “U-shaped” deformation patterns bridging sulcal banks coincide with the locations of short association fibers, supporting the idea that these pathways may arise passively from mechanical elongation rather than active axonal guidance alone (Garcia et al., 2021; Pron et al., 2021)

### 4.3 Implications for tractography and modeling

The present framework offers a novel route to geometry-informed tractography. Conventional tractography relies entirely on diffusion data, which is prone to crossing-fiber ambiguities and orientation noise (Schilling et al., 2019). In contrast, our method provides a deformation-derived orientation prior derived directly from cortical geometry. Such a prior could serve as an initialization or regularization term in existing probabilistic tractography algorithms - e.g., in complex tissue interfaces (Shastin et al., 2022; St-Onge et al., 2018) -, reducing spurious connections and improving anatomical plausibility. This idea is further corroborated by recent deep learning models show it is possible to approximate tractography using only T1w images (Cai et al., 2024; Tan et al., 2025).

Because the model requires only a structural MRI, it is applicable in populations where diffusion data are unavailable or unreliable, such as fetal or neonatal imaging. The approach also enables new avenues for personalized modeling: the deformation field captures an individual’s unique folding pattern and could be used to infer individualized fiber scaffolds or to generate subject-specific biophysical priors for connectome reconstruction. By extending to the whole cortex, our findings also align with prior work in the hippocampal domain, where unfolding the structure aided diffusion-based tractography (Hussain et al., 2020).

### 4.4 Limitations and future work

Several limitations should be noted. First, the current implementation treats folding as an approximately reversible process, omitting growth-induced remodeling or axonal plasticity that may occur during development. Including viscoelastic relaxation or differential growth between cortical layers could improve biological realism. Second, although the quasi-static model reproduces large-scale deformation patterns, finer microstructural features such as laminar tension and anisotropic material properties are not yet modeled explicitly. Third, validation was performed on a single longitudinal fetal subject and one adult; broader datasets will be required to establish generalizability across individuals and developmental stages.

The mesh quality significantly influences the simulation’s computational efficiency and precision, with suboptimal meshing leading to erroneous results.(Dogan & Atilgan, 2008; Osman et al., 2025). However, the computational demands of the method are modest: volumetric mesh generation required approximately 12 minutes, the unfolding simulation 20 minutes, and synthetic bundle generation *<*10 seconds per tract on a standard desktop CPU (Intel® Core™ i9-14900KF). These runtimes suggest that subject-specific geometry-derived orientation fields can be generated efficiently without specialized hardware, making the approach feasible for large-scale or developmental studies.

Future work will integrate the deformation-derived orientation fields with diffusion-based tractography, using the former as a prior to guide or regularize streamline propagation. Alternatively, one could apply the deformation to entire whole-brain tractograms to study developmental structural dynamics (Di Stefano et al., 2025; Karimi et al., 2025). This will allow direct quantitative comparison of geometry-informed and diffusion-informed connectivity maps. Previous studies indicate fiber development may involve stress-dependent growth along folding directions. The present framework elongates fibers in the direction of deformation but does not explicitly incorporate stress-dependent growth. Including such mechanisms could improve accuracy but would introduce additional parameters lacking well-defined biological counterparts. Limiting the method to structural MRI visualizes geometric information encoded in folding without relying on any diffusion-derived data. Extending the model to multi-layer cortical meshes or multimodal registration frameworks could further elucidate how folding mechanics influence functional network topology.

## 5 Conclusion

This work demonstrates that subject-specific cortical geometry alone can reveal major aspects of white-matter organization. By leveraging the quasi-static reversibility of cortical folding, we reconstructed individual deformation trajectories from structural MRI and derived corresponding white-matter orientation fields without using diffusion data. The resulting geometry-driven fiber orientations reproduced diffusion-derived patterns with high regional correspondence and produced biologically plausible strain distributions aligned with known zones of folding complexity and variability.

These findings establish cortical folding not merely as a surface feature but as a volumetric anatomical constraint that encodes the mechanical and geometric principles shaping white-matter architecture. The proposed framework provides a novel, anatomically grounded prior for tractography—one that can be applied even in the absence of diffusion imaging. Beyond tractography, the ability to infer deformation fields and local strain from morphology opens opportunities for studying the biomechanics of brain development, individual variability, and disease-related folding abnormalities.

Future developments will integrate this geometry-informed model with diffusion-based tractography and multimodal imaging, enabling more accurate, individualized representations of brain connectivity grounded in both anatomy and mechanics.

## Data and Code Availability

Each custom pipeline step, including meshing, XPBD simulation, fiber generation, and deformation, are implemented in the open-source tractography visualization tool NeuroTrace. (https://gitlab.com/Besm/neurotrace).

## Author Contributions

B.O. (data curation, writing, editing, experimental design, methodology, software tools); R.V. (supervision, editing); A.J. (conceptualization, supervision); K.G.S. (data curation, experimental design, writing, editing, conceptualization, resources); M.C. (experimental design, conceptualization, writing, editing, supervision);

## Acknowledgments

We thank Gauri Sharma for helping with proofreading and textual suggestions. This work was funded in part by the Dutch Research Council (NWO, OCENW.M.22.352) awarded to MC. KGS is supported by the National Institute of Health through award K01-EB032898. Data were provided by the developing Human Connectome Project, KCL-Imperial-Oxford Consortium funded by the European Research Council under the European Union Seventh Framework Programme (FP/2007-2013) / ERC Grant Agreement no. [319456]. We are grateful to the families who generously supported this trial.

## Ethics

No primary data were collected for this study. All analyses were performed on previously acquired MRI datasets from the Developing Human Connectome Project (dHCP) and the Human Connectome Project (HCP). The original studies involving human participants were reviewed and approved by their respective institutional ethics committees, and written informed consent was obtained.

## Supplementary Material

**Figure 1:**
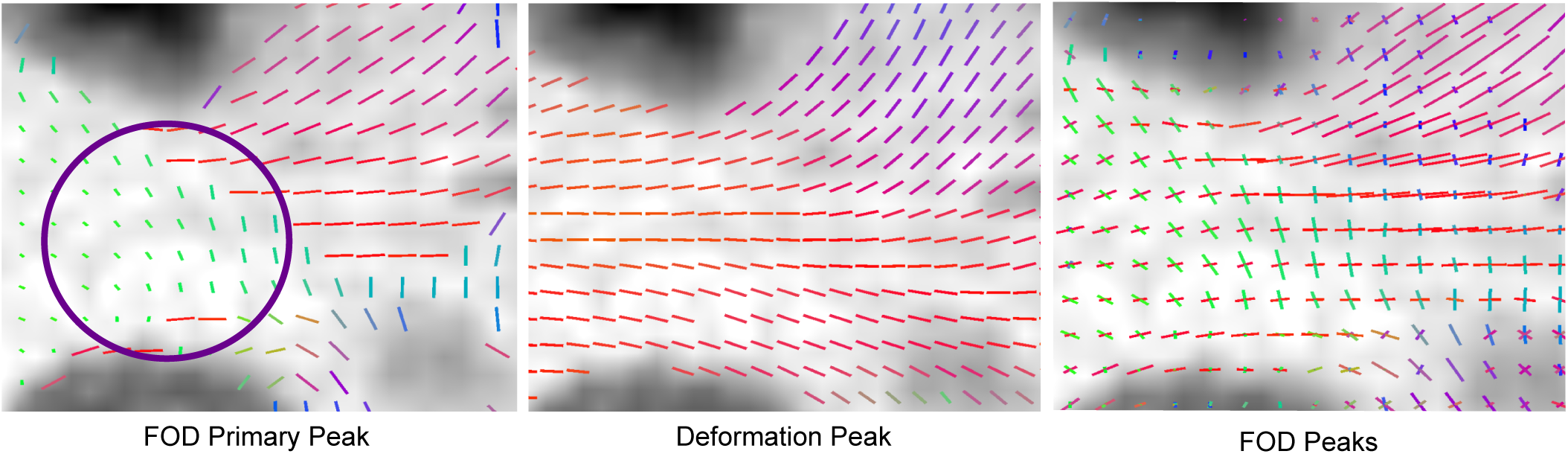
Region with low-scoring primary peak alignment (purple circle), where deformation-derived peaks align with secondary peaks.

